# Measuring Endoplasmic Reticulum Signal Sequences Translocation Efficiency Using the Xbp1 Arrest Peptide

**DOI:** 10.1101/281535

**Authors:** Theresa Kriegler, Anastasia Magoulopoulou, Rocio Amate Marchal, Tara Hessa

**Author notes:** To whom correspondence should be addressed: Phone: Int+46-8-16 24 60. Fax: Int+46-8-15 36 79.

## Abstract

Secretory proteins translocate across the mammalian ER membrane co-translationally via the ribosome-sec61 translocation channel complex. Signal sequences within the polypeptide, which guide this event, are diverse in their hydrophobicity, charge, length, and amino acid composition. Despite the known sequence diversity in the ER-targeting signals, it is generally assumed that they have a dominant role in determining co-translational targeting and translocation initiation process. We have analyzed co-translational events experienced by secretory proteins carrying efficient, versus inefficient (poorly hydrophobic) signal sequences, using an assay based on Xbp1 peptide-mediated translational arrest. With this method we were able to measure the functional efficiency of ER signal sequences. We show that an efficient signal sequence experiences a two-phases event in which the nascent chain is pulled from the ribosome during its translocation, thus resuming translation and yielding full-length products. Conversely, the inefficient signal sequence experiences a single weaker pulling event, suggesting inadequate engagement by the translocation machinery of these marginally hydrophobic signal sequences.

## Introduction

Proteins sorted via the secretory pathway are co-translationally translocated into the endoplasmic reticulum (ER) membrane. To select for secretory proteins, the cell scans for the presence of an N-terminal signal sequence or hydrophobic transmembrane domains (TMD) within the nascent polypeptide (Blobel and Dobberstein, 1975). When such a hydrophobic sequence on the nascent protein emerges from the ribosome, it binds the cytosolic signal recognition particle (SRP). This physical interaction defines the recognition stage of the ribosome/nascent chain targeting reaction (Walter and Blobel, 1981, Kurzchalia et al., 1986). Hydrophobic sequence-containing ribosome nascent chain complexes (RNC) are guided to the Sec61 translocation channel at the ER membrane via binding to the SRP receptor (Gilmore et al., 1982). Upon docking on Sec61 complex, the elongating polypeptide chain moves directly from the ribosome exit tunnel into the Sec61 translocation channel (Park and Rapoport, 2012). Signal sequences or TMD integrate into the membrane by the translocation channel via the lateral gate (Van den Berg et al., 2004, Voorhees and Hegde, 2016). Analysis of stalled RNCs of secretory proteins at different lengths shows signal sequences to crosslink to the lateral gate helices of the Sec61 complex (Jungnickel and Rapoport, 1995, Plath et al., 1998). Membrane integration by the translocation channel requires certain hydrophobicity to be able to partition into the lipid bilayer (Hessa et al., 2007, Van den Berg et al., 2004, Voorhees and Hegde, 2016). Simultaneously with this translational-coupled membrane integration, signal sequences invert to acquire N_in_-C_out_ orientation and, subsequently, their cleavage by the signal peptidase occurs at the luminal side of the ER membrane.

Although signal sequences show no homology in their primary structure, they have several common features. In general, they contain a positively charged amino-terminal region (N-domain), a hydrophobic core region (H-domain), and a polar carboxyl-terminal region, that usually contains the processing site for the signal peptidase (von Heijne, 1985). The signal sequence performs essentially two functions: it serves as a signal for targeting of ribosome-nascent chain complexes to the ER, and is necessary for translocation initiation across the ER membrane (Blobel and Dobberstein, 1975, Walter and Blobel, 1981). Several studies have shown that sequence variation among signal sequences can affect the efficiency of protein targeting, translocation, and signal sequence cleavage (Kim et al., 2002, Levine et al., 2005). Further, recognition of some signal sequences by the translocation channel requires the contribution of accessory factors, which also suggests that differences among signal sequences influence their interaction with the Sec61 protein complex (Fons et al., 2003, Jungnickel and Rapoport, 1995).

More recently, considerable attention has been paid to ribosomal mediated translational stalling as a method for measuring co-translational events such as folding and TMD insertion into the ER membrane (Ismail et al., 2012, Nilsson et al., 2015). Translational stalling occurs at specific amino acids in the nascent peptide during co-translational protein folding (Yanagitani et al., 2009, Yanagitani et al., 2011); one known example is represented by the mammalian X-box binding protein 1 (Xbp1) stalling sequence (Yanagitani et al., 2009). Recent work employing ribosome profiling has led to the identification of the potential critical amino acid responsible for the arrest in the Xbp1 sequence (Ingolia et al., 2011). Although the exact stalling mechanism remains yet to be identified, it has been shown that Xbp1 stalling sequences can be a useful tool to monitor co-translational translocation and insertion processes (Ismail et al., 2012).

In the present study, we exploit Xbp1-mediated ribosomal stalling to study the co-translational folding events: translocation, orientation, and cleavage of signal sequences during their insertion into the ER. We focused on measuring and characterizing the force exerted on signal sequences by using the arrest peptide derived from Xbp1 (Yanagitani et al., 2009). Our results revealed a two-phases pulling force profile when an efficient (hydrophobic) signal sequence was studied, in which the first-phase represents early engagement with the translocation channel and the second-phase represents inversion of the signal sequence. The force profile was highly dependent on the hydrophobicity of the H-domain and N-terminal charged residues for successful insertion, inversion, and cleavage of the signal sequence. In contrary, an inefficient (less hydrophobic) signal sequence experienced only a weak single-phase pulling force event, suggesting inefficient engagement and insertion. This might explain the functional efficiency of different signal sequences based on the pulling force exerted on them by translocation channel complexes during the early stage of translocation, and therefore determining the downstream consequences of the proteins localization.

## Results

### Experimental framework

Translational arrest peptides (APs) are short polypeptide sequences that interact with the ribosome exit tunnel and stall translation. Co-translational events such folding or membrane insertion can generate forces on the nascent chain able to break these interactions, therefore allowing translation to resume. These forces can be monitored by fusing the arrest peptide (force sensor) C-terminally to a nascent chain, and following its release from translational arrest. To measure the pulling force exerted on signal sequences we choose to use prolactin (Prl), a well-studied secretory protein that is efficiently targeted and translocated across the ER membrane. We started by identifying the frame in which the forces act on the emerging nascent chain by placing an Xbp1-based arrest-peptide, a 25 aa long mammalian peptide derived from the X-box binding protein 1, referred to as AP, (for more specific modifications see Table 1, STAR Methods) at different distances (L) downstream from the start methionine of Prl (Figure 1A and S1). We used several Xbp1 versions for different constructs, to allow for better dynamic range (Table 1, STAR Methods). For Prl, we used an version of Xbp1 that, in the absence of any pulling force, arrested translation of nascent chains by ∼70% (Figure 1B).

**Table 1:**
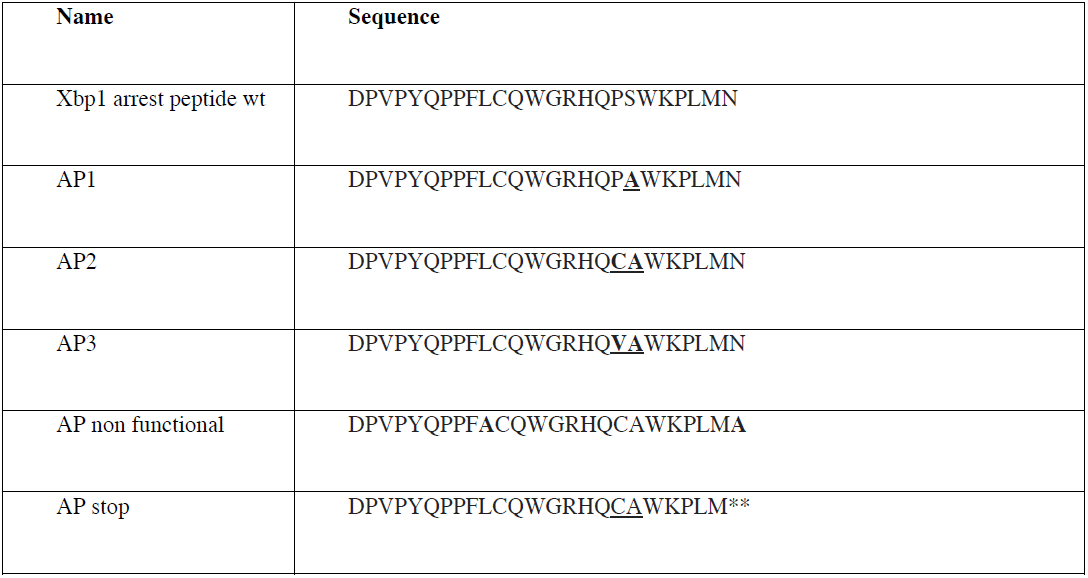
Different variants from the Xbp1 derived arrest peptide that were used in this study.

**Figure 1.**
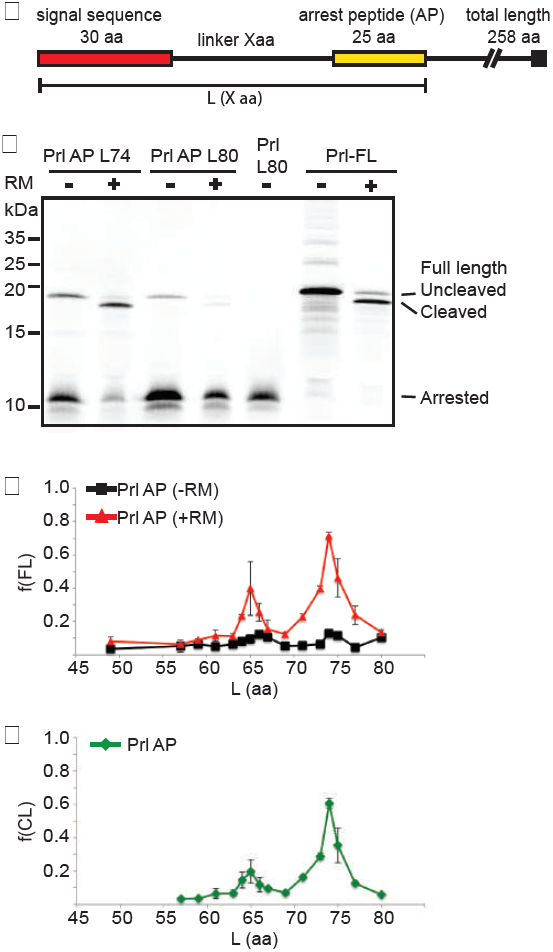
Xbp1 arrest peptide-mediated pulling forces of efficient signal sequence. **A.** Schematic overview of the model protein, prolactin (Prl) used for measuring the pulling force. The mammalian arrest peptide Xbp1 (AP) was placed at various distances from the N-terminus signal sequence of Prl (linker X aa). In the presence of pulling force acting on the nascent chain, the full-length protein is produced (774 aa), in absence of such a force the translation stalls at the arrest peptide and a shorter version of the protein is produced (arrested form, 110 aa). The distance between the start methionine and the C-terminal of the arrest peptide varies in each construct indicated as L (X aa). **B**. *In vitro* translation of Prl AP intermediates was synthesized in a rabbit reticulocyte lysate coupled transcription-translation reaction in the presence (+) or absence (-) of dog pancreas rough microsomes (RMs).[35S]-methionine labeled reactions were subjected to RNAse treatment and polypeptides were resolved by SDS-PAGE. The intensities of the protein band were quantified by phosphorimaging. Position of full-length released Prl, its precursor and arrested fraction is indicated. Apparent molecular masses are shown in (kDa). **C**. Pulling force profile of Prl AP. The *f*_*FL*_ for each truncated construct plotted as a function of L amino acids in presence (red) and absence (black) of RMs. Profiles represent at least three data points, the standard deviation of the data set is shown. **D**. Prl AP signal sequence cleavage profile plotted as function of the L. Fraction cleaved signal sequence (*f*_*CL*_), presented as the cleavage ratio of cleaved/(cleaved + uncleaved + arrested) from each reaction.

Upon co-translational translocation of Prl signal sequence via the Sec61 channel and folding in the ER lumen, nascent chains will experience a variety of tensions. If a strong pulling force acts on the chain at the moment of Xbp1-mediated arrest, translational stalling will be released, and hence the ratio between full-length and arrested protein will increase, as determined by SDS-PAGE. To test this, proteins were expressed in Rabbit Reticulocyte Lysate system in the absence or presence of rough microsomal membranes (RM). Both arrested and full-length proteins, of cleaved and un-cleaved species, were detected (Figure 1B). We calculated the full-length fractions (fraction _FL_) by quantifying both the cleaved and uncleaved population of translated products, and dividing the full length ones by the total products, including the arrested fractions.

### Efficient signal sequence generates a two-phase pulling force

To identify relevant different variants of expressed proteins, we created two different control mutants, one with a stop codon at the arrest site (Prl L80) to only obtain the constructs products migrating at the size of the arrested protein, and the other one with a non-functional arrest peptide (Prl FL) to only create the one migrating at the molecular weight of the full-length product. Figure 1B shows the expected size of the full-length protein and the arrested fragment. Note that we observed a small leakage (∼30%) in the system, where the arrest was overcome, generating full-length protein even in the absence of RM (Figure 1B).

Our data shows that two RM-dependent pulling events take place on nascent chains carrying an efficient signal sequence: one when the nascent chain is 65 residues (L65), where about 30 residues are exposed from the ribosome, and the second at L74, with about 39 residues exposed (Figure 1C). In the absence of ER membranes little pulling force was observed at the corresponding lengths, indicating that the main force exerted was related to interactions with the membrane and/or membrane components (Ismail et al., 2012). Signal sequence cleavage was quantified relatively to the total translation reaction, and the fraction of cleaved sequence, f(Cleaved), is represented in Figure. 1D, consistent with the pulling force profile for L=74, indicating that as the nascent chains are pulled from the ribosome they are accessed by the signal peptidase for cleavage. Less cleavage was noticed for constructs with a shorter linker length, L65, reflecting a pulling state in which the nascent chain was starting to be engaged by the Sec61 channel, but the signal sequence was not yet fully accessible to the signal peptidase in the membrane. We assumed in this case, that the fraction of cleaved protein observed was generated from the full-length protein spontaneously released in the absence of forces.

### Characterization of arrested signal sequence

To understand the translocation state of these intermediate lengths in their translationally, arrested versus pulled nascent chains, we subjected the membrane to Proteinase K digestion of intermediate lengths L57, L65, L74 and L80, representing the two pulling events and two arrested intermediates (Figure 2). The results show that the full-length uncleaved protein, in all cases, was accessible to PK, confirming that it represents RM-independent spontaneous release to the cytosol. By contrast, the cleaved full-length protein, which appears only in the presence of RM, was always PK-resistant, indicating that the RM-dependent pulled species were translocated to the ER lumen coincidently with signal sequence cleavage. In the presence of membranes, the released full-length cleaved proteins, protected in the lumen of the microsomes, were highest for L65 and L74, the most pulled intermediate. The arrested intermediates L57 and L80 instead, showed almost no protected cleaved fragments (Figure 2A). The arrested proteins showed a successive increase of arrest fragments, protected from PK as the chain length was extended, where the short intermediate L57 was completely digested, and the longer L80 seemed to be mostly protected, indicating that the longer arrested chains were stably associated with the membranes (Figure 2A and B).

**Figure 2.**
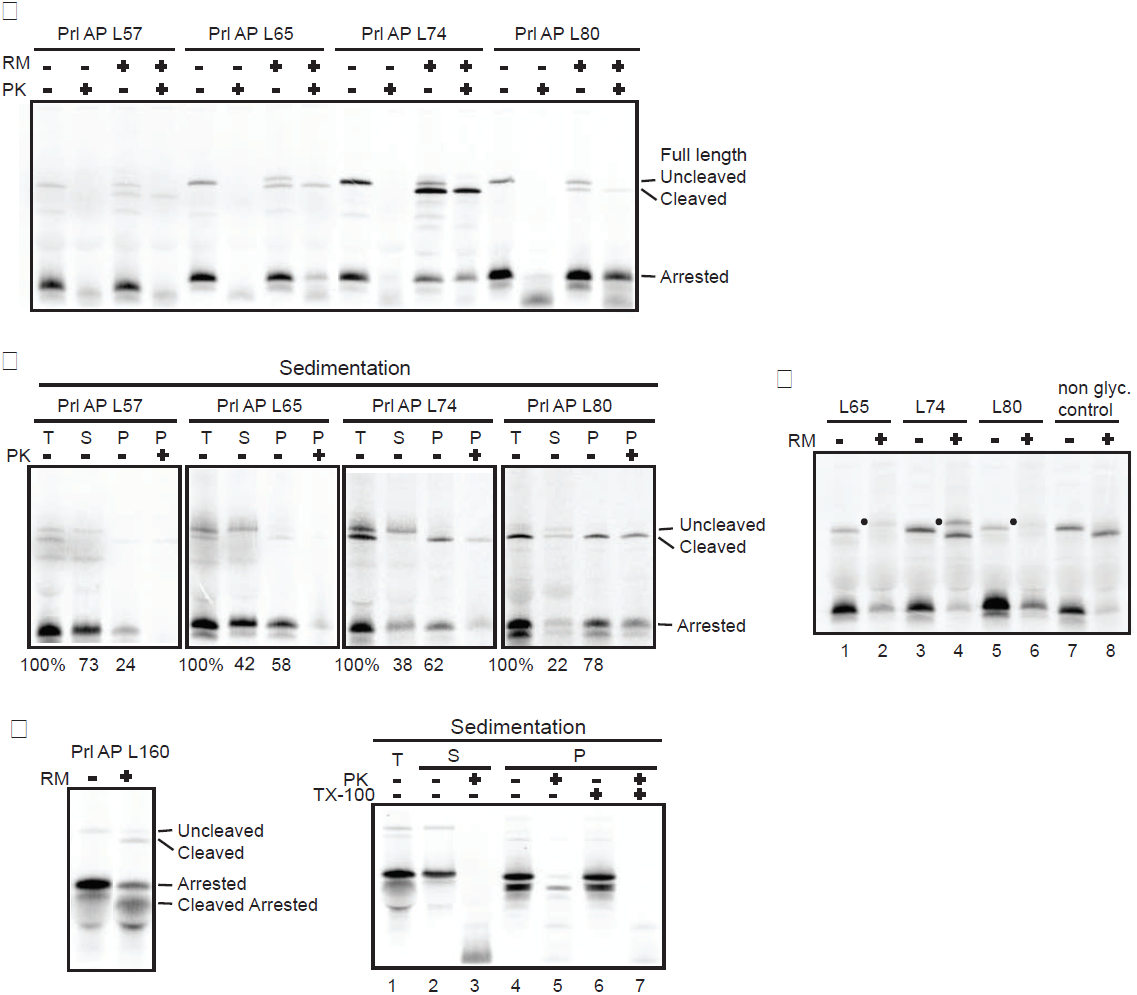
Analysis of Prl AP translocation efficiency across the ER membrane. **A.** Translocation of Prl AP (Xbp1) intermediates; L57, L65, L74 and L80 were examined by protease protection in presence and the absence of dog pancreas rough microsomes (RMs). No RM samples serves as control reaction. Equal aliquots of completed translations were subjected to proteinase K-digestion on ice for 1h and untreated samples are shown for each intermediate. Only the translocated proteins are protected after digestion of the translation reactions. The full-length protected proteins lanes 4, 8, 12 and 16 and arrest protected proteins lanes 8, 12 and 16 are shown. All reactions were RNase treated to release the nascent chain from the ribosome followed by resolving on SDS-PAGE. **B**. *In vitro* expression of intermediates in **A** was subjected to sedimentation on sucrose cushion under physiological salt conditions for the intermediates L57 and L65. Sedimentation yields membrane (P) and supernatant (S) fractions. The pellet fraction was split into two equal volumes, were one was analyzed directly (-PK) and the other one treated with proteinase K (+PK). The precursor and mature protein fractions are indicated (un-cleaved and cleaved) and the stalled nascent chain on the ribosome indicated as arrested. Percent membrane bound (P) and non-bound (SN) respectively are shown. **C**. Prl AP L65, L74 and L80 intermediates carrying a single glycosylation site 38 amino acids downstream of the arrest peptide were expressed in the absence and presence of RMs. Only translocated polypeptides into the lumen side will be modified and give rise to 2 KDa higher molecular weight band (•). Cleaved signal sequence and the mature glycosylated protein are shown (lane 2, 4 and 6). **D**. *In vitro* translation of Prl AP L160 intermediates showing the arrested Prl and the cleaved arrested form in the presence of membrane (left panel). Positions of full-length released Prl, its precursor, and the arrested cleaved fraction are marked by arrows. After sedimentation of Prl AP L160 intermediate in the presence of RMs, the pellet fraction was split into four equal volumes, were one was treated with PK (lane 5), one treated with Triton X-100 (TX-100) (lane 7) and the other two no treatment serves as controls.

To further assess whether the translation intermediates were membrane bound and in an engaged translocation state, or free in the cytosol, we performed sedimentation experiments at physiological salt concentrations, to separate translation mixtures into membrane bound pellet and a cytosolic supernatant. As expected, the cleaved released FL proteins, protected from PK digestion, segregated into the pellet fraction, confirming their translocation in the ER lumen. The un-cleaved released FL proteins were instead detected in the soluble fraction. Figure 2B also shows that the most pulled intermediate lengths L65 and L74, were 58 % and 62% respectively (similarly under high salt condition, Figure S2) of the arrested chains, and were associated with membrane but not protected from PK digestion, indicating that most of the stalled chains were associated with membrane but not protected from PK digestion. Interestingly, the least pulled intermediates L57 were mostly in the soluble fraction, whereas the majority of L80 chains were associated with the membranes, indicating that shorter translationally arrested RNCs, representing early translocation intermediates, are less stably associated with the membrane, while the longer ones are in a more engaged state and remain tightly bound to the membrane. RNCs carrying Prl AP L57 represent an early translocation intermediate; although the signal sequence can likely bind to SRP, it is too short to interact productively with the translocation channel (Jungnickel and Rapoport, 1995). Therefore, the RNCs bind to the ER membrane only loosely, and the transition to tight binding does not take place. Prl AP L65 represents a stage were the nascent chains starts to engage fully with the translocation channel. At this stage the chain gets longer, and likely to be more membrane associated and engaged, hence to increase the pulling effect exerted on the chain.

We also assessed whether these translocation intermediates had bound to RMs in a salt-resistant manner. Several previous studies have shown that upon initial transfer to the translocation channel, nascent chains are bound in a salt-sensitive manner (Jungnickel and Rapoport, 1995, Zheng and Nicchitta, 1999). Only after additional elongation (after approximately 60 total amino acids) the nascent chain is bound in a salt-resistant manner that is dependent on a functional signal sequence. Analysis of Prl AP L57 to L80 translocation intermediates of the signals revealed that, with the exception of L57, all of the constructs were bound to the membrane in a salt-resistant manner (Figure 2B and S2).

To further confirm the translocation state and cleavage of the signal sequences, we introduced a glycosylation site into the mature part of the Prl AP lengths L65, L74 and L80 as an indicator of protein translocation and exposure to ER lumen. We observed that Prl AP chain lengths L74 were glycosylated, and that the signal sequence was cleaved (Figure 2C, lane 4), indicating that RM-dependent pulled population was translocated to the ER lumen. Both Prl AP L65 and Prl AP L80 were glycosylated to a lesser extent (Figure 2C, lane 2, 6), suggesting that only a small fraction of these intermediate chains were able to reach the luminal side of the ER, consistent with the signal sequence cleavage data.

Next, we generated longer intermediates of L110 and L160 in order to understand whether, and/when, the signal sequence of the arrested population was cleaved. Recent study shows the signal sequence of Prl truncated nascent chains to be processed between lengths 137 and 163 amino acids (Devaraneni et al., 2011). We observed that, at L110, the majority of the nascent chains were still in an arrested uncleaved state similar to L80 (data not shown), while at L160 we observed the arrested RNC population signal sequence to be cleaved (Figure 2D, left panel). Separation of membrane bound arrested fractions, in the presence of PK treatment, showed the cleaved signal sequence to be protected, and that this protection was abolished upon membrane solubilization by detergent (Figure 2D, right panel and S2). In summary, our results show that the nascent chains at L65 and L74 underwent a conformational change, and that, under an arrest peptide driven elongation arrested state, they were pulled out from their arrest. The first-phase pulling event is presumably representing an early engagement stage with the Sec61 translocation channel, where very little cleavage occurs and the arrested RNCs appear to be partially membrane-bound and protected. The L74 intermediate appears to represent a second stage, where signal sequence is pulled and undergoes inversion during its transitioning from the Sec61 channel to the lipid bilayer for efficient cleavage (Jungnickel and Rapoport, 1995, Martoglio et al., 1995, Mackinnon et al., 2014, Voorhees and Hegde, 2016).

To gain more insight into when translocation of the Xbp1 arrest occurs, we investigated the timing of translocation and signal sequence cleavage of the pulled and arrested intermediates of Prl AP at lengths L65, L74, L80 and L160 (Figure S3A-D). At intermediate length L65 we observed a time lag of 5min between pulling and signal sequence cleavage, indicating that polypeptide chains were pulled first and then cleaved (Figure S3A). Interestingly, translocation intermediate length L74 showed that the pulling of the nascent chain increased linearly with signal sequence cleavage (Figure S3B). The arrested population, on the other hand, showed a greater time lag between the small pulled fraction and signal sequence cleavage (Figure S3C). For the longer intermediates L160, although the majority of the nascent chains (> 90% in total) were in an arrested state quite early on, we noticed the arrested fraction signal sequence to be up to 60% cleaved after 15 min translation initiation (Figure S3D). In conclusion, time course measurements showed that pulling events preceded signal sequence cleavage for the L65 intermediate, whereas the L74 intermediate pulling and cleavage seemed to occur simultaneously, suggesting that signal sequence at this chain length is optimally positioned to get cleaved by the signal peptide peptidase. This supports the idea that the first pulling peak at L65 represents the early engagement stage, evident by the time delay between pulling and cleavage, and the L74 intermediate represents the second stage with the pulling and cleavage occurring simultaneously.

### Positive charges in the N-region of the signal sequence promote pulling

Next we examined the potential role of the N-region in the pulling events. We generated a Prl signal sequence version where all the charged and polar residues were substituted with Ala residues, keeping the residues in the vicinity of the cleavage site intact (Prl 5LAP) (Figure 3A). We measured the pulling force for these constructs at different lengths in the presence and absence of microsomes. To our surprise the second pulling event was abolished upon omitting the charged residues in the N-region of the Prl signal sequence, and the full-length Prl protein was only recovered without any signal sequence cleavage. A potential explanation is that the second pulling event might occur when the signal sequence undergoes inversion into the correct orientation required for cleavage by the peptidase. We also observed a higher first pulling event compare to the WT Prl signal sequence, probably due to the more hydrophobic nature of the Prl 5L sequence a stronger pulling is experienced. To test whether the charges are responsible for the inversion of the signal sequence, we re-introduced 2 Lys residues into the n-region generating (Prl 5LKKAP), and indeed we were able to restore the second pulling peak that resulted in the signal sequence to be cleaved (Figure 3B), suggesting the importance of n-region charges for the second pulling event to occur.

**Figure 3.**
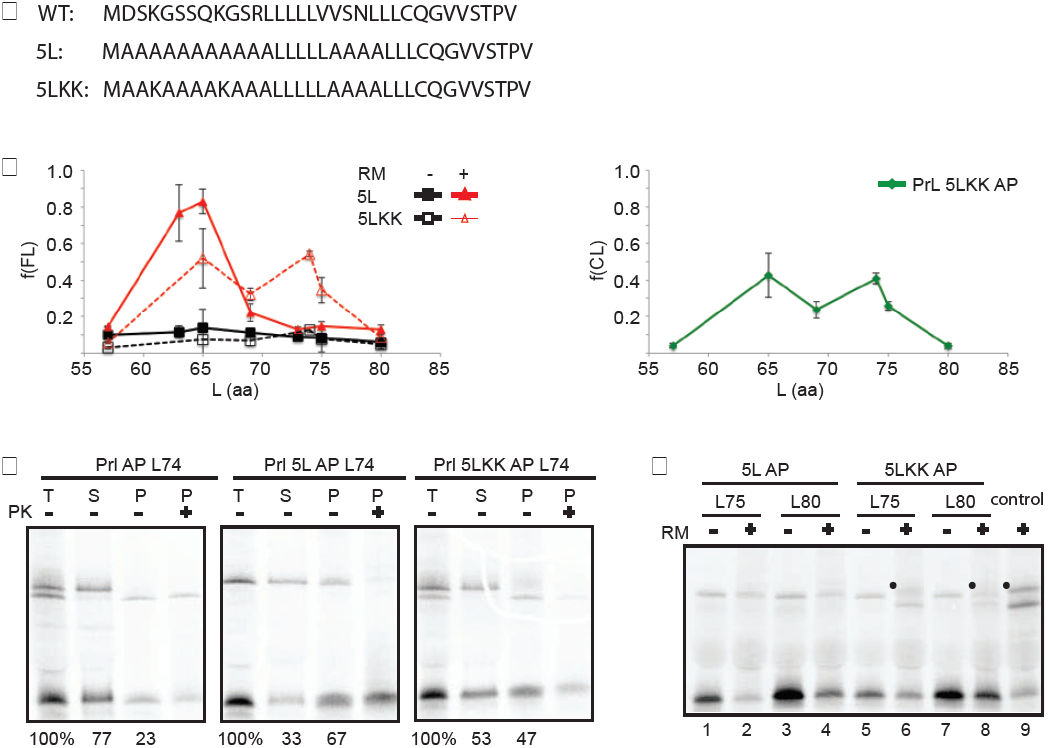
Pulling force is dependent on the signal sequences composition. **A**. Sequence of the wild type and mutant Prl signal sequences 5L and 5LKK, hydrophobic core and N-terminal region are modified indicated in bold type and Lysine mutant underlined. **B**. Quantifications of fraction full-length protein of Prl 5L AP and Prl 5LKK AP plotted as function of L amino acids for constructs shown in A in presence (red) and absence (black) of membrane (left panel). Fraction cleaved signal sequence (mature protein) of Prl 5LKK AP in presence of membrane (green) (right panel). **C**. *In vitro* expressed constructs; Prl AP, Prl 5L AP and Prl 5LKK AP of intermediate length L74 was sedimented though a high salt sucrose cushion, pellet fractions were split into two samples and one sample was subjected to PK digestion. Numbers represent efficiency of protein translocation in percent of the mature form (data are from three experiments, the standard deviation of the data set is shown). **D**. Translocation efficiency of Prl 5L AP and Prl 5LKK AP constructs with linker lengths L75 and L80 assessed using diagnostic glycosylation site (•). Control experiment with efficient glycosylation is shown.

We then examined whether the arrested Prl 5LAP and Prl 5LKKAP were partitioned in the membrane and/or translocated into the ER-lumen, by performing high-salt sedimentation experiments. Unlike with the wild-type signal sequence, where the uncleaved full-length represents a cytosolic spontaneous release, the un-cleaved full-length Prl 5LAP was partly sedimented in the membrane pellet fraction. The sedimented full length however was completely digested upon PK treatment (Figure 3C), suggesting the possibility that Prl 5LAP was associated with the lipid bilayer on the cytosolic side of the membrane. In contrast, the Prl 5LKKAP constructs had cleaved signal sequences, and were cleaved and protected from digestion consistent with the restored pulling force profiles (Figure 3C). We further examined the translocation state by using the diagnostic glycosylation assay where we tested the intermediates L75 and L80. Both Prl 5L AP intermediate lengths appeared not to be glycosylated, in contrast to the Prl 5LKKAP L75 and L80 intermediates were modified, confirming its correct translocation across the membrane (Figure 3D). These results clearly demonstrate that the second pulling event, at chain length L74, likely representing a step where signal sequence is in a more integrated state where it can undergo inversion and cleavage. Further this also highlights the importance of N-region charges of the signal sequences for its translocation and correct orientation in the membrane.

In addition, our data is in agreement with the recent cryo-EM structure of the mammalian ribosome-Sec61 translocation channel complex, where the N-terminal signal sequence of 86 amino-acid long prolactin was inserted head-in, intercalated between the lateral gate helices within the channel (Voorhees and Hegde, 2016, Goder and Spiess, 2003). The signal sequence is subsequently being inverted, resulting in the C-terminus in the lumen ready for cleavage by the signal sequence peptidase on the luminal side of ER (Goder and Spiess, 2003, Devaraneni et al., 2011).

### The effect of hydrophobicity on the signal sequence pulling

Hydrophobicity of signal sequence has been established as an important determinant that influences the efficiency of targeting and translocation of secretory proteins (von Heijne, 1985). We wondered whether by increasing the hydrophobicity of the H-region, hence improving the efficiency of the signal sequence, we would detect any changes in the pulling force. To this end, we designed a more hydrophobic version of Prl signal sequence containing 12 Leu in the H-region (Prl 12LAP), while we kept the rest of the sequence intact as the WT including the N-region charges (Figure 4A). Similarly, multiple linker-length constructs were generated and tested for their pulling ability. The results showed one dominant pulling event for Prl 12LAP at length L61 and a second smaller pulling at length L71-L73 (Figure 4A). Compare to the WT Prl signal sequence pulling force profile, we observed a higher first-phase pulling event and a notable reduction of the second-phase event (Figure 4B). Earlier cross-linking study showed that RNCs of nascent chain length 50-60 residues were too short to interact productively with the translocation channel (Jungnickel and Rapoport, 1995). Therefore, we subjected translated 12LAP L61 nascent chains to salt-resistant sedimentation assay and accessibility to proteinase K. Our data showed the majority of the pulled nascent chains, that were cleaved and PK protected, segregate into the pellet fraction (Figure 4C). We also confirmed that 12LAP L57, 61 and 65 were all translocated across the membrane by probing them with diagnostic glycosylation site (data not shown). Additionally, we noticed that the arrested chains were partially membrane associated but sensitive to PK-digestion, indicating that the arrested were not tightly bound to the membrane as well as judged by the amount of arrested chains in the cytosol (Figure 4C). These results demonstrate that by increasing hydrophobicity of the signals sequence we were able to improve the early engagement step (or insertion), representing the first-phase pulling event. This in turn improve the subsequent inversion of the signal sequence, hence the efficient cleavage of 12L intermediate is evident.

**Figure 4.**
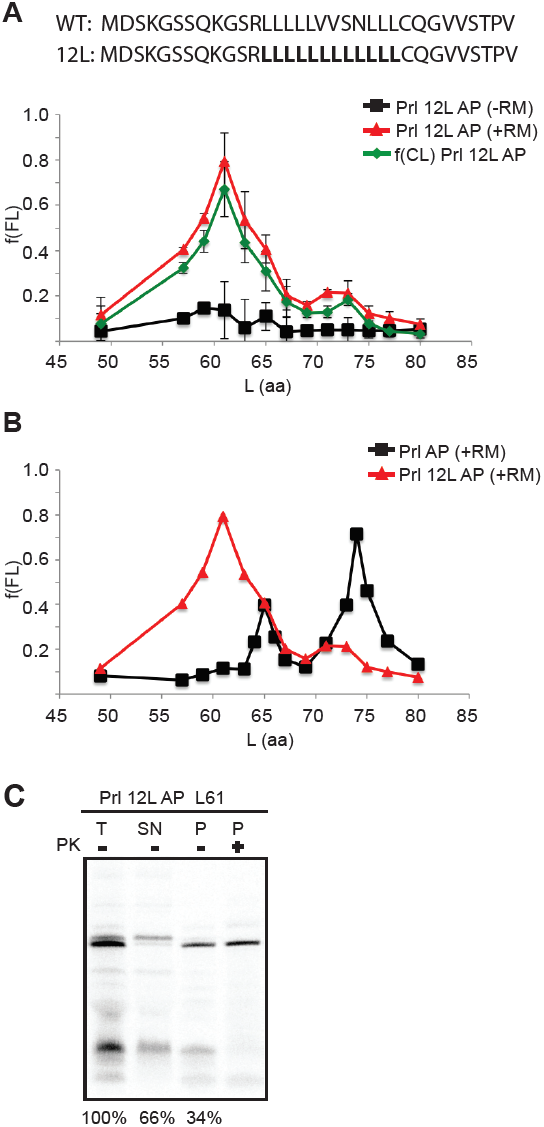
Hydrophobic core of a signal sequence is primary determinant of pulling force. **A**. Sequence of mutant Prl signal sequences Prl 12L AP, hydrophobic core are modified indicated in bold type. **B**. Quantifications of fraction full-length protein (*f*_*FL*_) of Prl 12L AP plotted as function of L amino acids for the construct shown in A in presence (red) and absence (black) of membrane. In addition, fraction cleaved signal sequence polypeptides (green) are shown. **C**. Sedimentation experiment combined with PK-digestion of Prl 12L AP with linker length L61. **D**. Comparison of wild-type Prl AP and Prl 12L AP pulling force profiles.

### Force profiles of inefficient signal sequences

Next, we investigated the pulling force profiles acting on an inefficient signal sequence (Kim et al., 2002, Hessa et al., 2011). Earlier studies showing non-equivalence among signal sequences (von Heijne, 1985) and their requirements for certain factors (Fons et al., 2003) prompted us to test whether pulling force were exerted on these inefficient signal sequences. As a test protein we used the signal sequence of prion protein (PrP), a mammalian glycoprotein, mainly expressed and anchored to the cell surface by a glycophosphatidylinositol (GPI) linkage. Topologically, PrP contains an N-terminal signal sequence, a marginally hydrophobic transmembrane domain (TMD), and a foldable C-terminal unit of three helical domains before the GPI attachment site (Prusiner et al., 1998).The PrP signal has been proved to less effectively interact with the Sec61 translocation channel, resulting in PrP adopting three different topologies and/or escaping from the translocation channel altogether into the cytosol (Hegde et al., 1998, Hessa et al., 2011).

We substituted the signal sequence of Prl with PrP signal sequences (PrP-Prl). Constructs carrying inefficient signal sequence were tested as described before, by placing the Xbp1 at various distances from the PrP signal sequence (Figure 5A). The criteria for the choice of Xbp1 version is based on the capability of the chain to be pulled from the ribosome PTC site at minimum of 50% and maximum 100% arrest fraction in order to measure a detectable force. We initially used two other Xbp1 mutant versions to scan for the pulling capability of arrested nascent chains, but we were unable to capture any significant pulling event with these variants (Table1, STAR Methods and Figure S4). We turned to a third Xbp1 mutant version (PrP-Prl AP) (Table1, STAR Methods) that it has been shown to be able to arrest a hydrophobic transmembrane domain consisting of 6L/13A sequence by 50% (Schiller N. et al., 2017).

**Figure 5.**
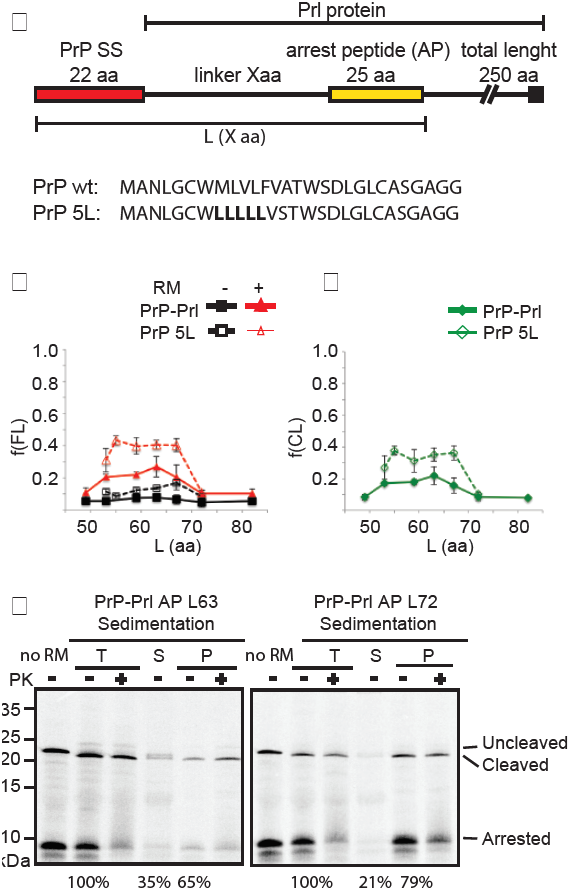
XBP1 arrest peptide-mediated pulling forces of inefficient signal sequence. **A**. A schematic of the construct containing prion protein (PrP) signal sequence (25amino acids) fused to the C-terminal of the Prl model protein (PrP-Prl). A weaker version of the mammalian arrest peptide Xbp1 was used at various linker lengths (linker Xaa) positions from the start methionine to generate translational stalling at different length (L in aa). **B**. Force profiles obtained for the constructs PrP AP and PrP 5L AP respectively. In the presence of RMs (red curves) and absence (black) of membrane. **C**. Fraction cleaved signal sequence (*f*_*CL*_) for constructs in b plotted as function of the L. Data represent at least three experiments; the standard deviation of the data set is shown. **D**. The efficiency of translocation was determined by centrifugation of translated PrP-Prl linker lengths L63 and L72 in to yield supernatant (S) and pellet (P). Fractions cleaved, uncleaved and arrested are indicated. Numbers represent efficiency of protein translocation in percent of the mature form (data are from three experiments, the standard deviation of the data set is shown).

The force profiles for PrP-Prl AP showed a less evident pulling event ranging from linker length L53 to L67 corresponding to the H-region in the PrP signals sequence (Figure 5B). Although signal sequence of the full-length, pulled population appeared to be efficiently cleaved (Figure 5C), the pulling peak was lower compared to Prl signals sequence pulling profile. By increasing the hydrophobicity of the signal sequence (PrP5L-Prl AP) the pulling force in the presence of membrane was higher, but it was observed also in the absence of membrane indicating that the sequence context surrounding the hydrophobic core may be as important as hydrophobicity for signal sequence insertion (Figure 5B and 5C). Therefore, we replaced the hydrophobic core of Prl signal sequence with the one of PrP to generate the chimera Prl-PrPAP, consisting of the Prl charged N-region, the PrP hydrophobic-region and C-terminal cleavage site region of Prl. The pulling force profile of the chimera resembled the WT Prl force profile with typical two-phase pulling event, but less efficiently pulled (Figure S5A). To confirm the specific pulling effect detected for PrP-PrlAP constructs, we substituted Prl-signal sequence with a 25 unstructured Gly-Ser residues, as expected no pulling force could be detected for GS sequence at various lengths (Figure S5B). In addition, sedimentation analysis under physiological salt conditions combined with PK digestion of L63 (pulled) and L72 (arrested) intermediates showed that 65% and 79% respectively were sedimented with RM. We also observed that only small percentage of the FL released and arrested chains in the cytosolic fraction (Figure 5D). Overall the pulling force profile and sedimentation experiments suggested that PrP-PrlAP RNCs were stably associated with the membrane. When we precipitated the membrane fraction using CTABr we noticed that membrane-bound nascent chains in the pellet fraction of PrP-Prl AP L72 and Prl AP L80 RNCs were equally associated with the ribosomes (data not shown).

Taken together, our results showed that less pulling force was exerted on the PrP-signal sequence, and that the sequence signatures, including the N-terminal charges and the hydrophobicity, play a major role in determining if a stalled nascent chain preceded by an inefficient signal sequence could be pulled by the translocation channel and/or by any other accessory factors in the membrane. These results are supported by earlier studies on inefficient signal sequences being associated with different translocation channel complex (Fons et al., 2003, Conti et al., 2015)

### Arrested nascent chains interacts with the Sec61 complex

To determine the immediate interacting environment at the membrane for the arrested Prl nascent chains of various linker lengths, we used bis-maleimidohexane (BMH) that specifically crosslinks accessible cysteines in the nascent chain to the proximate neighboring proteins. Translation of Prl AP L57-L80 was carried out in the presence of ER membranes, and membrane-bound nascent chains were isolated on sucrose cushion (to asses only the membrane bound nascent chains) and subjected to immunoprecipitation using specific antibody against Sec61 channel components. BMH crosslinking of Prl AP linker lengths L57-L80 revealed that the arrested fractions were associated mainly with two products of sizes ∼10 and ∼40 KDa corresponding to Sec61α and Sec61β, with maximum intensity crosslink to L74 and L80 (Figure 6A and S6A). Interestingly, when similar analysis was performed on the Prl12LAP L61-L80, we observed that the main crosslink was to Sec61β, appearing at L63 trough L75 (strongest at L69), but not to the Sec61α at any intermediate length (Figure 6B). Since the majority (∼80%) of the polypeptide (L61) in this case was pulled, we were unable to capture any interactions between this small arrested population and Sec61α, indicating that Prl 12LAP being pulled by the Sec61complex with a faster kinetics at the indicated intermediate length, and as the nascent chain proceeds in length (L69-L75) the arrested 12L remained associated with Sec61β (Figure S6B). Analysis of BMH crosslinking for PrP-Prl AP at linker lengths L63 and L72 (most pulled versus arrested states) gave similar crosslinks bands corresponding to Sec61α and Sec61β starting at the shortest intermediate L63, and increasing with the higher arrested fractions (L72 and L82) (Figure 6C, and S6C). This suggested that this early and persistent interaction with the Sec61 complex could be due to the PrP signal sequence recognition by Sec61 complex and transition to the membrane occurred with a slower kinetics. Further, immunoprecipitation against Sec62 antibody resulted in no particular crosslink to PrP-Prl AP at any lengths (L49-L80) (data not shown), consistent with previous studies showing this interaction with Sec62 and Sec63 to occur beyond nascent chain length of 130 amino acids (Devaraneni et al., 2011).

**Figure 6.**
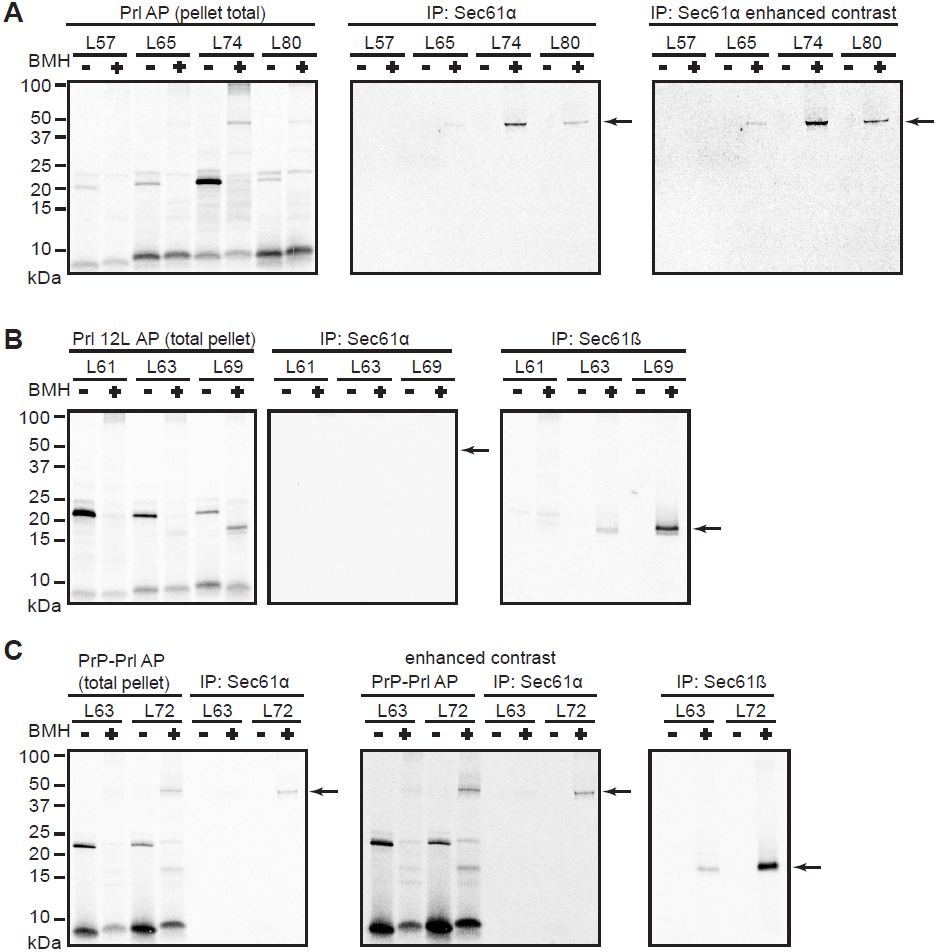
Arrested nascent chains interactions with the Sec61 complex. Autoradiograph of radiolabelled translation products of nascent chains containing various signal sequences at the indicated intermediate lengths treated with the chemical cross-linker BMH in the presence of RMs and sedimented on sucrose cushion. Pellet fractions were analyzed by SDS-PAGE directly or after being subjected to immunoprecipitation against anti-Sec61α and β antibodies (see also Figure S6A-C). **A**. The position of crosslink products in the presence and absence of BMH are indicated (arrows) for Prl AP intermediates total pellet and immunoprecipitation (IP) with Sec61α and Sec61β (see Figure S6) are shown. **B** and **C.** As in A, total pellet and immunoprecipitation (IP) against anti-Sec61α and β antibodies for Prl 12L AP and PrP-Prl AP intermediates respectively.

All together these results clearly demonstrated that the arrested Prl nascent chains were in immediate contact with the sec61 complex, indicating that nascent chains were pulled from their arrested state via the Sec61 translocation channel (Jungnickel and Rapoport, 1995, Voorhees and Hegde, 2016). When the signal sequence was converted to more hydrophobic one, the transition to the membrane seemed to occur faster, therefore decreasing the contact time with the Sec61 components; on the other hand, this transition was delayed for the inefficient signal sequence (PrP-Prl).

## Discussion

In the present study, we describe new insights into co-translational processes experienced by nascent signal peptides during translocation at the ER membrane using Xbp1-mediated translational arrest. By using stalled ribosome nascent chains, derived from the arrest peptide of Xbp1, we were able to measure the functional efficiency of ER signal sequences. Insertion and processing of signal sequences generates a force that stretches the nascent chain in a most favorable conformation around the PTC site, in order for the next aminoacyl-tRNA to form a peptidic bond and for the elongation to resume, thus overcoming the arrest. This force is reflecting any co-translational interaction sensed by the ribosomal tunnel/translocation channel/membrane lipids during nascent chain progression. Here we compare the pulling force generated by highly efficient versus inefficient signal sequences by examining nascent chains of different length, representing snapshots of the biogenesis of secretory proteins, during co-translational folding. We have found a two-phase pulling force exerted on the efficiently targeted and translocated signal sequence of prolactin at chain lengths 65 and 74 amino acids. The first pulling event is representing an early engagement stage with the Sec61 translocation channel, where the signal is minimally cleaved and the arrested RNCs appear to be partially membrane-bound and protected. The second pulling event, at L74 intermediate, appears to represent a second stage, in which signal sequence is integrated into the Sec61α as shown by the recent structure, where the signal of the prolactin nascent chain is intercalated between the lateral gate helices, and undergoes to a subsequent inversion and cleavage in the lipid bilayer (Voorhees et al. 2016, Jungnickel and Rapoport 1995, Martgolio et al. 1995). Further, our data shows that signal sequence cleavage for the arrested population occurs at chain length L160 in agreement with recent study (Devaraneni et al. Cell 2011), (Figure 2D). Our crosslinking analysis supports the idea that the initial interaction between Sec61complex and Prl signal sequence occurs at L65 intermediate, as shown by a weak crosslink to Sec61α and β, and that this interaction is optimal for the L74 intermediate, when nascent chains undergo through the second pulling event (Figure 6A). These data is in agreement with previous studies showing that RNCs, at these particular lengths (L65 and L74), are efficiently cross-linked to the Sec61 translocation (Jungnickel and Rapoport, 1995, Mackinnon et al., 2014).

Signal sequence mutants (5L-Prl) lacking charges in the N-region produce only one pulling force peak while the second is abolished. Such construct does not integrate fully into the membrane, fails cleavage and glycosylation unless charged residues are reintroduced into the N-region (5LKK-Prl) consequently restoring all the mentioned modifications (Figure 4A and B). This confirms the “head-in-first” model, with the N-terminal signal sequence being inserted in toward the ER lumen, and subsequently inverted resulting in the C-terminus in the lumen ready for cleavage by the signal sequence peptidase on the luminal side of ER (Goder and Spiess, 2003, Devaraneni et al., 2011, Voorhees and Hegde, 2016). This highlights the importance of the positive charged residues of the N-region for the inversion process to happen. Our data additionally show that when the hydrophobicity of the H-region is increased the pulling force is shifted towards a strong earlier pulling (Prl12L, length intermediate L61), with fully cleaved signal sequence, implicating that the majority of the chains undergoes a faster transition into the membrane, when sequence content is optimal. Indeed, we only observed crosslinking of 12L to the Sec61 β protein suggesting that the signal sequences were pulled via the Sec61 complex with a faster kinetic.

Our data obtained with the signal sequence derived from prion protein show a lower degree pulling experienced by the nascent chain, reflecting the sequence inefficiency, poor hydrophobicity, and the lack of charged residues at the N-terminal. Given the stable association with the membrane and efficient crosslinking to the Sec61 complex, these differences in the signal sequence result in prolonged sensing by the Sec61 channel at the tested intermediate lengths. In conclusion, the signature features here described for efficient signal sequences appear to be critical and required for effective pulling of the nascent chain from the ribosome.

Furthermore, our data provide evidence of “hydrophobicity barrier” at the ribosome-Sec61 translocation channel-membrane, where a certain hydrophobicity threshold has to be overcome in order for the signal sequence to be pulled and transitioned into the membrane lipids via lateral gating (Hessa et al., 2007, Hessa et al., 2005). However, an inefficient signal sequence (PrP) is unable to overcome such barrier completely, evident from the less pulling exerted of the stalled chains and the prolonged association with the Sec61 complex, indicating that its complete transition the into membrane might dependent on additional factors. Accordingly, recent studies showed for the PrP signal sequence cleavage to be delayed, and to be associated with a sub-complex of Sec61, Sec62 Sec63 (Conti et al., 2015). Further work will be needed to determine the impact of these associated sub-complexes (Sec62/63) on pulling force of these moderately hydrophobic signal sequences.

## Material and Methods

### Enzymes and Chemicals

All Enzymes used in these studies were from Thermo Scientific. The oligonucleotides for cloning and PCR reactions were obtained from Eurofins Genomics. [^35^S]-methionine was from Perkin Elmer. All chemicals used were from Sigma-Aldrich unless differently specified. Plasmid DNA, RNA and for gel extraction kits were from Thermo Fisher and were used accordingly to the manufacturer’s protocols.

### Plasmids and Constructs

All constructs were sub-cloned into pSPBP4 expression vector, with an inserted 950-base-pair fragment encoding bovine pre-prolactin, described previously (Krieg, Walter and Johnson, 1986). We used a site-directed mutagenesis approach to introduce a SpeI restriction site upstream of the initial methionine of the prolactin signal sequence, and a KpnI restriction site downstream of the signal sequence. The 25 amino acids long arrest peptide from the X-box binding protein (Xbp1) (Yanagitani et al., 2011) was introduced into Prl-SpeI-KpnI at position 85, from the start methionine or at position 135, generating a construct of total length of 110 or 160 amino acids until the arrest, respectively. The Xbp1 arrest peptide was introduced in the constructs by Gibson assembly method (Gibson et al., 2009) using the oligomers in Table 4. All variants of Xbp1 arrest peptide showed in Table 1 were created by site directed mutagenesis. Shorter length than 85 amino acids between the start and the arrest peptide were obtained by PCR using forward and revers primers that were complementary to the regions indicated in Table 3 and in Figure S1 with right-facing and left-facing arrows, respectively. Glycosylation site (NST) was introduced by site directed mutagenesis 36 amino acids after the C-terminal of the arrest peptide. The prion protein (PrP) signal sequences – prolactin (Prl) chimera, was generated by inserting PrP signal sequence in between the SpeI and KpnI sites (Table 2) by using 2-step annealing oligomers method (Hessa *et al.*, 2005). Shorter constructs were generated as before using the same truncation primers (Table3). When ligation strategy was used, phosphorylation of the oligonucleotides was performed using the T4 Polynucleotide Kinase from Thermo Fisher, according to manufacturer’s protocol. All constructs were transformed into chemically competent DH5-α cells.

**Table 2:**
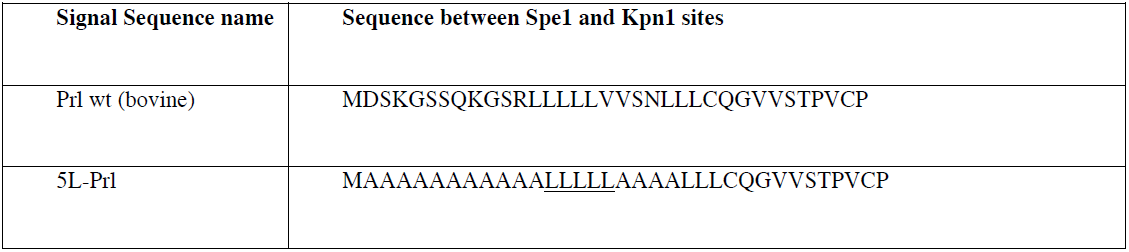
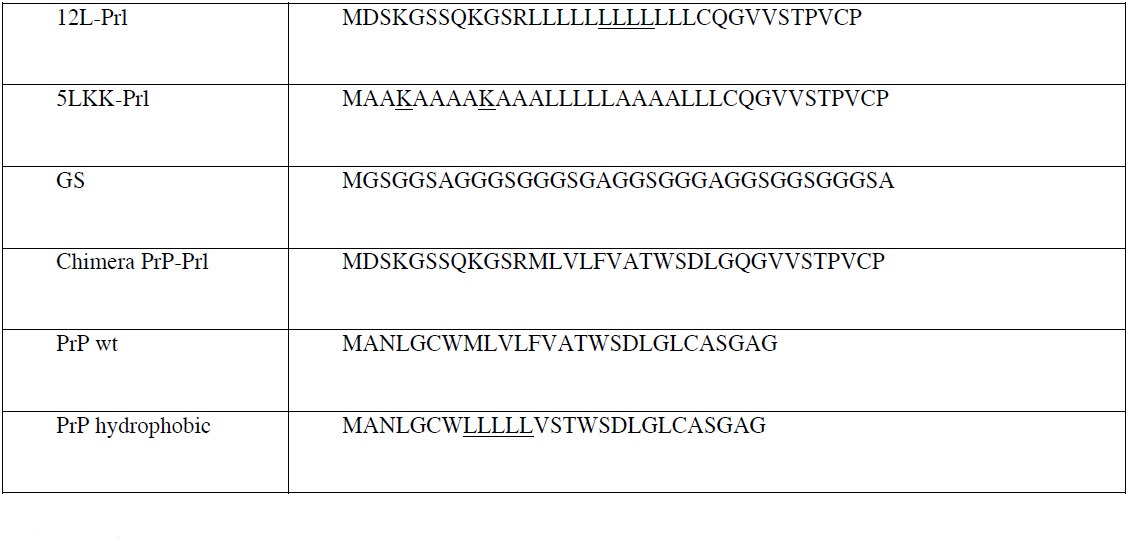
Different Signals Sequence variants that were analyzed in the pre-prolactin background in this study.

### In vitro transcription/translation

All transcription-translation reactions were performed using a Rabbit Reticulocyte Lysate (RRL) derived SP6 coupled system (Promega) for 30 min at 30 °C in the presence of 2.5 μCi [^35^S]-methionine in a total volume of 10 μl. Reactions were performed with addition of 0.5 μl column washed microsomes (RM) from dog pancreas (tRNA products) when indicated. Proteinase K digestions were for 1 hour on ice. The reactions were stopped by addition of sample buffer and boiling for 5 min at 95 °C before treatment with RNaseA (15 min, 37 °C). Samples were separated by SDS-PAGE on 12% Tris-glycine polyacrylamide gels and analyzed by autoradiography.

Additional experiments were performed using an uncoupled system (Sharma et al., 2010) starting from purified PCR products. Transcription was performed with an SP6 polymerase (source) for 1 h at 40 °C. Transcripts were then added to the translation reaction containing 2.5 μCi [^35^S]-methionine in 10 μl reactions and 0.5 μl RMs if indicated. Sample preparation, RNase treatment, visualization and analysis were as described above.

### Sedimentation on sucrose cushion

The selected constructs were incubated in RRL for 30 min at 30 °C. Samples (20 μl total volume) were loaded on a 80 μl sucrose cushion (low salt: 250 mM sucrose, 100 mM KAc, 50 mM Hepes pH 7.9, 5 mM MgAc; high salt: 250 mM sucrose, 500 mM KAc, 50 mM Hepes pH 7.9, 5 mM MgAc) and were centrifuged for 4 min at 46 000 rpm at 4°C in a Optima MAX-XP Ultracentrifuge with a TLA 100.3 rotor (Beckman). The pellet was suspended in (20 μl) sucrose suspension buffer (50 mM Hepes 7.9, 250 mM sucrose, 100 mM KAc, 5 mM MgAc) and analyzed directly or treated with proteinase K for 1 hour on ice (Gorlich and Rapoport, 1993). All samples were resolved by SDS-PAGE and analyzed as described above.

### CTABr precipitation

Precipitation of the Peptidyl-tRNA from cell-free translations with cetyltrimethylammonium bromide (CTABr) was performed as previously described (Gilmore and Blobel, 1985). In brief, translation products (40 μl) were added to 250 μl of 2% (w/v) CTABr. Precipitation of peptidyl tRNA was induced by the addition of 250 μ1of 0.5 M sodium acetate (pH 5). The CTABr precipitate was incubated for 10 min at 30°C, the precipitates were then collected by centrifugation and washed twice with 500 μl of acetone:HCI (19:1) to remove CTABr. The supernatant was treated with 10% trichloroacetic acid (TCA) to recover protein contents and analyzed similarly. Samples then dissolved in SDS containing sample buffer for polyacrylamide gel electrophoresis analysis.

### Bis-maleimidohexane crosslinking and immunoprecipitation

Transcription/translation reactions were performed as described above, diluted in 10 volumes salt buffer (20 mM HEPES pH 7.4, 100 mM KAc, 5 mM MgCl_2_) and treated with 250 μM 1,6-bis-maleimidohexane (BMH) in DMSO for 30 min on ice. Control fractions were treated with DMSO. The reactions were quenched by adding 25 mM β-mercaptoehtanol and incubating for 10 min on ice. Samples were then loaded on an equal volume of sucrose cushion and the membrane fraction isolated as described above. Pellets were suspended in denaturing buffer (1% SDS, 0.1 M Tris pH 8), heated for 5 min at 95°C. Products were either analyzed directly by SDS-PAGE, or immunoprecipitated by dilution in10 volumes of cold IP-Buffer (100mM KAC, 50mM Hepes pH 7.4, 1% T-X-100). Anti-Sec61α, anti-Sec61β, anti-Sec62 or control anti-GFP antibodies were added, and samples were rotated over night at 4°C for anti-Sec61α and anti-Sec62 or 1.5 hours for anti-Sec61β and anti-GFP. After the primary incubation, protein A sepharose (Bio-Rad) was added to the samples followed by additional 2 hours of incubation at 4°C, rotating. The beads were washed three times with cold IP buffer and proteins eluted with 2x Sample buffer at 95°C for 5 min. Samples were analyzed by SDS-PAGE and autoradiography.

## Quantification and Statistical Analysis

Autoradiography images were acquired using a FLA-9000 phosphorimager equipped with ImageReader version 1.0 software (Fujifilm Corporation) and band intensities were quantified using Image Gauge software V4.23 (Fujifilm Corporation). Analysis of quantified bands was performed using Easy Quant (in-house developed quantification software), followed by normalizing the fractions to the number of methionine. Fraction of full-length protein (f(FL)) was calculated, as previously shown, by using the equation:

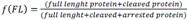

All data was confirmed in at least three (n=3) independent experiments (if not indicated differently in the figure legends). The error bars shown in all graphs represent the standard deviation between the different experiments as calculated by using Microsoft Excel. No additional statistical tests were performed.

